# Auditory evoked-potential abnormalities in a mouse model of 22q11.2 Deletion Syndrome and their interactions with hearing impairment

**DOI:** 10.1101/2023.10.04.560916

**Authors:** Chen Lu, Jennifer F. Linden

**Affiliations:** Ear Institute, University College London, London, WC1X 8EE, UK; Department of Neuroscience, Physiology, & Pharmacology, University College London, London WC1E 6BT, UK

**Keywords:** 22q11.2DS, Df1/+ mouse, hearing loss, auditory evoked potential

## Abstract

The 22q11.2 deletion is a risk factor for multiple psychiatric disorders including schizophrenia and also increases vulnerability to middle-ear problems that can cause hearing impairment. Up to 60% of deletion carriers experience hearing impairment and ∼30% develop schizophrenia in adulthood. It is not known if these risks interact. Here we used the *Df1/+* mouse model of the 22q11.2 deletion to investigate how hearing impairment might interact with increased genetic vulnerability to psychiatric disease to affect brain function. We measured brain function using cortical auditory evoked potentials (AEPs), which are commonly measured non-invasively in humans. After identifying one of the simplest and best-validated methods for AEP measurement in mice from the diversity of previous approaches, we measured peripheral hearing sensitivity and cortical AEPs in *Df1/+* mice and their WT littermates. We exploited large inter-individual variation in hearing ability among *Df1/+* mice to distinguish effects of genetic background from effects of hearing impairment. Central auditory gain and adaptation were quantified by comparing brainstem activity and cortical AEPs and by analyzing the growth of cortical AEPs with increasing sound level or inter-tone interval duration. We found distinctive measures of central auditory gain or adaptation that were abnormal in *Df1/+* mice regardless of hearing impairment, and other measures that were abnormal only in *Df1/+* mice with or without hearing impairment. Our data identify potential biomarkers for auditory brain dysfunction in psychiatric disease and illustrate that central auditory abnormalities in 22q11.2DS are a function of both genotype and hearing phenotype.

## Introduction

22q11.2 Deletion Syndrome (22q11.2DS) is the most frequent chromosomal microdeletion syndrome [1] and one of the strongest cytogenetic risk factors for a broad spectrum of psychiatric disorders [2,3]. For example, nearly 30% of 22q11.2 deletion carriers develop schizophrenia during their lifetime [4–6], and identified symptoms are indistinguishable between schizophrenia patients with the 22q11.2 deletion and those with idiopathic schizophrenia [5,7]. Insights gained from studying brain abnormalities in 22q11.2DS might therefore have general application to understanding brain abnormalities arising from vulnerability to psychiatric diseases such as schizophrenia. Mouse models of 22q11.2 deletion are also well established, enabling researchers to investigate, at the mechanistic level, what brain abnormalities arise from this genetic risk factor for psychiatric disorders and how experiential risk factors might affect their development [8].

One experiential risk factor for psychiatric disease that may be particularly relevant for 22q11.2DS patients is hearing impairment. Hearing impairment is a recognized experiential risk factor for schizophrenia in the general population [9,10]. Up to 60% of 22q11.2 deletion carriers have mild to moderate hearing impairment, primarily arising from chronic middle ear inflammation [11,12]. The possible contribution of hearing impairment to development of psychiatric disease in 22q11.2 carriers has not been studied. However, biomarkers of auditory brain dysfunction, such as abnormalities in cortical auditory evoked potentials (AEPs), are well-documented in 22q11.2DS (and schizophrenia) patients [13,14]. Surprisingly, the influence of hearing impairment on cortical AEP abnormalities in 22q11.2DS patients has not been systematically examined, despite the prevalence and inter-individual variability of hearing impairment in this population.

Here we investigated whether cortical AEP abnormalities in the *Df1/+* mouse model of 22q11.2DS depend only upon genotype or are affected by hearing phenotype. *Df1/*+ mice have a 1.2 Mb microdeletion homologous to the minimal 1.5 Mb deletion in 22q11.2DS [15] and replicate multiple physiological abnormalities observed in 22q11.2DS patients [16–19], including high inter-individual variation in hearing sensitivity. Like 22q11.2DS patients, *Df1/+* mice are susceptible to chronic middle-ear problems, and up to 60% have mild to moderate hearing impairment in one or both ears [20,21]. Moreover, previous work has suggested that auditory brain abnormalities in *Df1/+* mice correlate with hearing impairment [22]. Density of parvalbumin-immunoreactive inhibitory interneurons in the auditory cortex was abnormally low in *Df1/+* mice with hearing impairment, and gain of click-evoked cortical AEPs was abnormally high. No significant differences in AEPs were observed between *Df1/+* mice with normal hearing and their WT littermates; however, conclusions were limited by the use of a broadband click stimulus at a single sound level and repetition rate.

In the current study, we first conducted a comprehensive review of methods for cortical AEP measurement in mice, which revealed striking diversity in approaches. Adopting one of the simplest and best-validated strategies, we then investigated cortical AEP abnormalities in *Df1/+* mice in detail using pure tones, varying both tone intensity and inter-tone intervals to study gain and adaptation of auditory brain responses. Our data reveal that some cortical AEP abnormalities in *Df1/+* mice are independent of hearing impairment and others depend on degree of hearing impairment. The results point to specific AEP measures that could be reliable biomarkers for auditory brain abnormalities in 22q11.2DS patients, robust to individual differences in degree of hearing impairment.

## Methods and Materials

See Supplementary Information for additional details on acoustic stimulation, experimental procedures, and data pre-processing.

### Animals

Experiments were conducted in 29 *Df1/*+ mice (18 females, 11 males; mean age, 10.3 ± 1.4 weeks) and 22 WT littermates (8 females, 14 males; mean age, 10.2 ± 1.3 weeks). Sample sizes were chosen to be comparable to those in previous related studies [22]. *Df1/*+ mice were originally developed from a 129SvEvBrd X C57BL/6J background [15] and had been maintained on a C57BL/6J background for well over 25 generations, through pairing of *Df1/+* males either with WT females from the colony or (at least yearly) with newly acquired C57BL/6J females from Charles River UK. Mice were raised in standard cages and mouse housing facilities, on a standard 12 h-light/12 h-dark cycle. All experiments were performed in accordance with a Home Office project licence approved under the United Kingdom Animal Scientific Procedures Act of 1986.

### Recording procedures and measures

Electroencephalographic recordings were performed by an experimenter blind to the genotype of the animal, using procedures similar to those described previously [22]. Briefly, ketamine/medetomidine-anesthetized mice were oriented with the tested ear directed toward the speaker. The opposite ear was blocked with an earplug during recordings to ensure monaural stimulation. We recorded the auditory brainstem response (ABR) differentially from subdermal electrodes at the bulla of the tested ear and the vertex, and the cortical auditory evoked potential (AEP) single-ended from a subdermal electrode over the auditory cortex contralateral to the stimulated ear. The ABR is a sound-evoked signal that occurs within a few milliseconds of stimulus onset and consists of 5 successive waves arising from afferent activity in the auditory nerve and brainstem. The AEP is a later sound-evoked signal consisting of waves thought to originate from the thalamocortical projection (P1), the primary auditory cortex (N1), and higher auditory cortical areas (P2).

We used ABR and AEP recordings to obtain objective measures of hearing threshold, afferent auditory input to the brain, and contralateral cortical responses to auditory input for each ear in each mouse. More specifically, we used click-evoked ABRs to define hearing threshold; amplitude of tone-evoked ABR wave 1 (which arises in the auditory nerve) to quantify afferent input to the auditory brain; amplitudes and latencies of tone-evoked AEP waves to quantify auditory cortical activity; and comparative measures of AEP and/or ABR wave 1 amplitude to measure central auditory gain and adaptation (see Data analysis, below).

### Auditory stimuli

#### Click-evoked ABRs

Stimuli were 50 μs monophasic clicks ranging in sound level from 20 to 90 dB SPL in 5 dB steps, repeated 500 times at each sound level with an inter-onset interval of 50 ms.

#### Tone-evoked ABRs and AEPs

Stimuli were 16 kHz tones (5 ms, cosine-gated ramps) at 80 dB SPL, repeated 1000 times at an inter-tone interval (ITI) of 300 ms.

#### Level-dependent AEP growth functions

Stimuli were 16 kHz tones (5 ms, cosine-gated ramps) at 70, 80, 90, or 100 dB SPL, repeated 1000 times for each sound level with an ITI of 300 ms.

#### Time-dependent AEP growth functions

Stimuli were 16 kHz tones (5 ms, cosine-gated ramps) at 80 dB SPL, repeated 1000 times at ITIs of 200, 250, 300, 350 or 450 ms.

### Data analysis

ABR signals were filtered with a 100-3000 Hz band-pass filter before subtraction (vertex minus bulla electrode) to obtain the differential ABR signal. *Hearing threshold* was defined as the lowest click intensity level eliciting a characteristic ABR wave deflection at least twice as large as the time-dependent standard error in the mean click-evoked ABR waveform. *Tone-evoked ABR wave 1 amplitude and latency* were defined as the amplitude and latency of the peak of ABR wave 1 relative to baseline at stimulus onset; this peak was identified manually for each ear and animal from average ABR waveforms evoked by 16 kHz 80 dB SPL tones.

Three key deflections of the AEP waveform (P1, N1, and P2) were manually selected from the averaged waveform after removing heartbeat noise [23]. As in previous work [22], P1 was defined as the highest deflection between 15 to 30 ms post stimulus onset; N1 as the lowest deflection between 25 to 60 ms post stimulus onset; and P2 as the highest deflection between 60 to 120 ms post stimulus onset. Amplitude differences P1-N1 or N1-P2 were analyzed to minimize effects of baseline fluctuations on AEP measures. We quantified gain and adaptation of AEPs using three measures. *Central auditory gain* was defined as the ratio between the P1-N1 or N1-P2 AEP amplitude difference and the ABR wave 1 amplitude evoked by the same 16 kHz tone stimulation; this measure quantifies central auditory amplification of peripheral auditory nerve input. *Level-dependent AEP (LDAEP) slope* was defined as the slope of the best-fit linear function relating P1-N1 or N1-P2 AEP amplitude difference (or P1, N1 or P2 latency) to the sound intensity level; this alternate measure of gain was used to quantify central auditory excitability. *Time-dependent AEP (TDAEP) slope* was defined similarly as the slope of the best-fit linear function relating P1-N1 or N1-P2 AEP amplitude difference (or P1, N1 or P2 latency) to the logarithm of ITI duration; this logarithmic measure was used to quantify AEP adaptation to stimulus repetition.

The data were statistically analyzed and graphically visualized using Python. Generalized linear models (GLMs) were used to identify factors predictive of different outcome measures under a Gaussian distribution assumption (predictor variables: genotype; gender; age; hearing threshold of contralateral ear, and hearing threshold of ipsilateral ear). Further statistical analyses for each outcome measure were conducted using non-parametric statistical methods (randomization test, Spearman’s rank correlation test, Mann-Whitney U rank test, Kruskal-Wallis H test with post-hoc Dunn’s test). All significance tests were conducted two-tailed with α = 0.05, and exact p-values are stated where significant but greater than 0.001.

## Results

### Comprehensive literature review reveals a need for standardized approaches to auditory-evoked potential measurement in mice

Procedures for cortical AEP recording are relatively well established for human studies, but there is no standardized AEP recording paradigm for mice. To find out how AEP measurement in mice has been performed in the past, we reviewed 115 papers that were obtained from Web of Science using search terms ‘auditory evoked potentials’, ‘event-related potentials’, ‘mouse’, ‘rodent’. Papers that only covered auditory brainstem responses (sometimes called ‘early AEP’) were excluded. After consolidating the search results to eliminate papers using repeated methods, we still had 50 publications describing different approaches to AEP recording in mice. There was surprisingly little consistency between approaches used by unaffiliated labs. Electrode locations varied from study to study; some used single or symmetric electrode recording from one brain region and others recorded from multiple brain areas simultaneously (Figure 1A). Clustering all positive electrode locations across studies, we found that in the majority of previous studies, positive electrodes had been placed above auditory cortex, hippocampus and/or frontal areas (Figure 1B), while reference electrodes were typically located either at the occipital bone or over the frontal/olfactory lobe.

**Figure 1.**
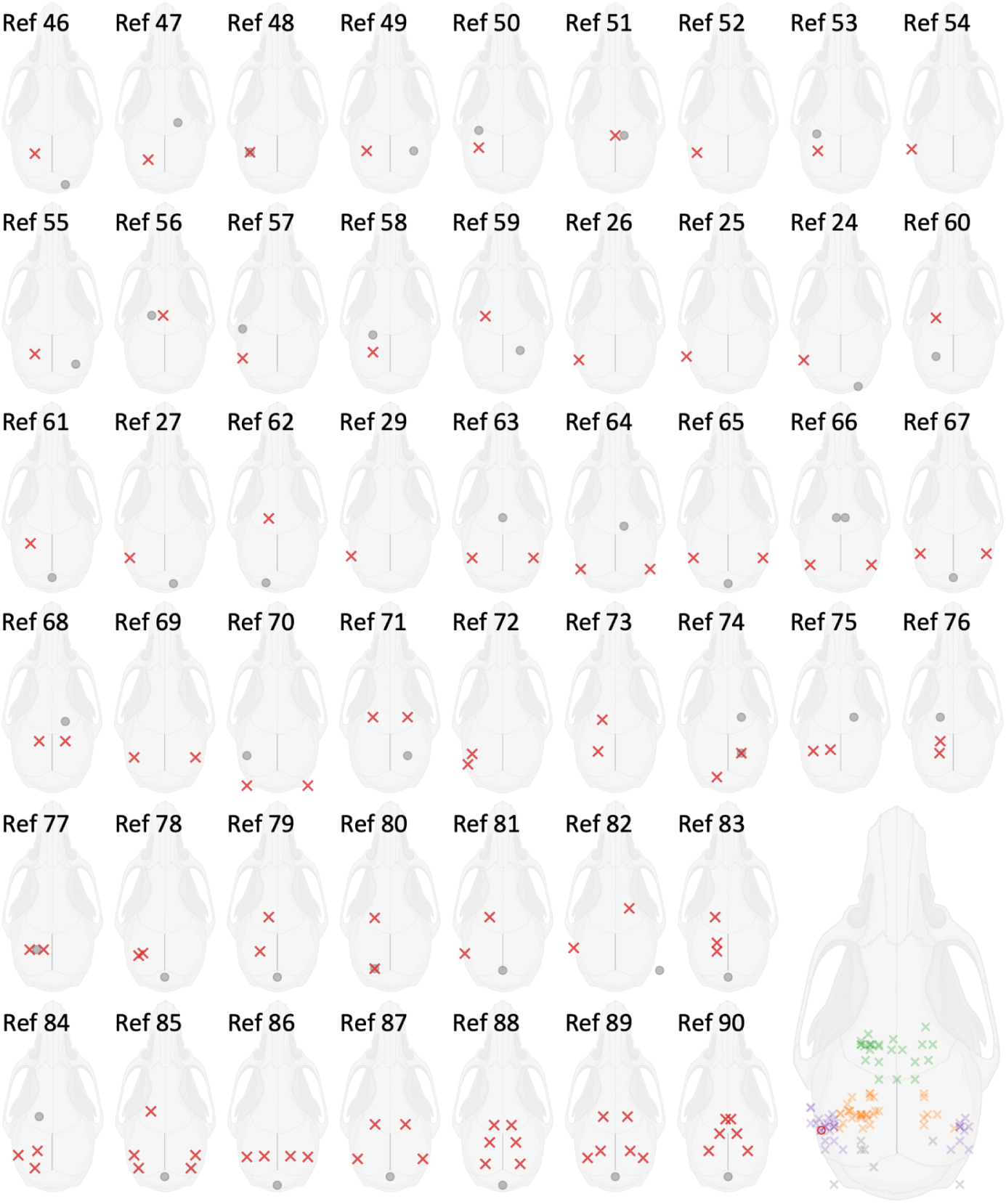
Comprehensive review of electrode placement strategies used for measurement of auditory evoked potentials (AEPs) in the mouse. (A) Electrode positions (red cross, positive electrode; gray circle, negative electrode) used for mouse AEP measurements in 50 publications representative of the diversity of approaches in 115 relevant studies identified through a literature search. (B) Summary diagram illustrating range of positive electrode positions used in previous studies. Positive electrodes were most frequently positioned over the auditory cortex (purple), frontal cortex (green), or hippocampus (orange).

In the absence of a standardized approach to mouse AEP recording, we adopted the simplest strategy compatible with other measurements of intracranial auditory cortical activity correlating with the AEP. Specifically, we placed a single subdermal positive electrode above the auditory cortex contralateral to the monaurally stimulated ear, and a reference electrode over the frontal/olfactory lobe [24–27]. Our definitions of the AEP wave deflections P1, N1 and P2 are consistent with definitions used in those studies [28]. We note also that the same electrode positioning with reversed polarity was used in one of the most detailed characterizations of subcutaneous cortical AEPs in mice to date [29]. Aside from the reversal of polarity, our AEP waveforms were consistent with those recorded by Postal et al. 2022, and our P1, N1 and P2 correspond to N23, P35 and N61 from their study.

We used this simple and well-validated approach to AEP measurement in mice to quantify AEP abnormalities in the *Df1/+* mouse model of 22q11.2DS, for comparison with AEP abnormalities observed in 22q11.2DS patients [30]. We focused particularly on a question that has been largely overlooked in previous studies of 22q11.2DS patients and 22q11.2DS model mice: are AEP abnormalities in *Df1/+* mice related to hearing impairment, a common comorbidity of 22q11.2DS?

### High inter-individual and inter-ear variation in hearing sensitivity in *Df1*/+ mice necessitates ear-by-ear analysis of effects of hearing impairment

We first confirmed previous findings that *Df1/+* mice replicate the high inter-individual and inter-ear variation in hearing sensitivity also observed in 22q11.2DS patients [12,20,22]. We quantified peripheral hearing sensitivity of each ear in each mouse using the click-evoked ABR (see Methods and Materials). ABRs were measured with subdermal electrodes (Figure 2A) during presentations of click stimuli varying in sound level from 20dB to 90dB SPL. The hearing threshold of an ear was defined as the lowest click intensity level eliciting a significant ABR wave deflection (Figure 2B).

**Figure 2.**
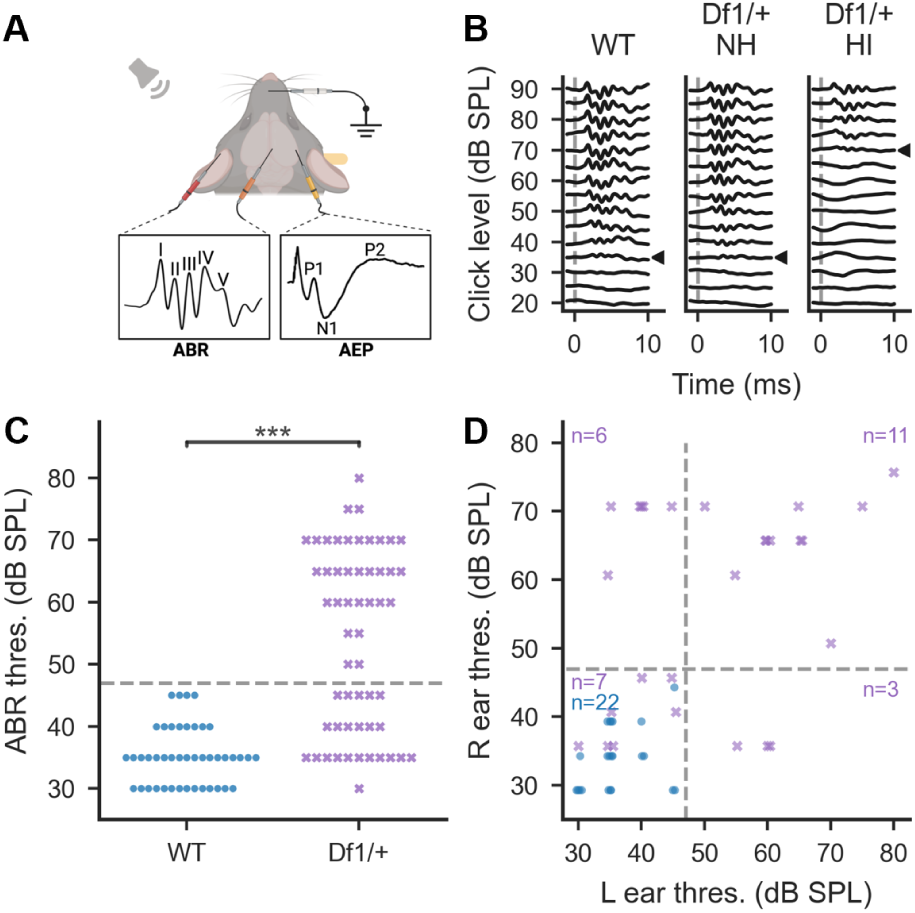
Experimental setup and ear-by-ear quantification of peripheral hearing sensitivity. (A) ABRs and AEPs were measured with subdermal electrodes in anesthetized mice. (B) Hearing sensitivity of each ear was quantified by identifying the click-evoked ABR threshold, i.e. the minimum click intensity eliciting a detectable ABR waveform. Three examples are shown, with click-evoked ABR thresholds indicated by arrowheads: left (WT) 35 dB SPL, middle (*Df1/*+) 35 dB SPL, right (*Df1/*+) 70 dB SPL. Dashed vertical line indicates click onset. (C) Distributions of click-evoked ABR thresholds in WT ears (blue circles) and *Df1/*+ ears (purple crosses). Ears with hearing loss were defined as those with click-evoked ABR thresholds more than 2.5 standard deviations above the mean WT threshold (i.e., above 47 dB SPL; dashed line). (D) Comparison of click-evoked ABR thresholds in the right and left ears; conventions as in C.

Hearing thresholds of *Df1/*+ ears were bimodally distributed (Figure 2C), as demonstrated previously [20,22]. On average, hearing thresholds for *Df1/*+ ears were significantly elevated compared to those from their WT littermates (*Df1/*+, 54.02 ± 14.41 dB SPL from 56 ears, WT, 35.23 ± 4.64 dB SPL from 44 ears; Wilcoxon rank-sum test, p < 0.001), but many *Df1*/+ ears had WT-like hearing thresholds. Here, we defined an ear as exhibiting hearing impairment if its hearing threshold was more than 2.5 standard deviations above the mean hearing threshold for WT ears (47 dB SPL). Then we grouped *Df1/*+ ears into “normal hearing” (*Df1/*+ NH, 23 ears) and “hearing impaired” (*Df1/*+ HI, 33 ears).

Notably, hearing impairment in one ear of a *Df1/+* mouse did not predict hearing impairment in the other. *Df1/+* mice often had hearing impairment in only one ear, and there was no significant correlation between the hearing thresholds of the two ears in *Df1/+* mice (Spearman’s rank correlation, ρ = 0.31, p = 0.12). Among all *Df1/*+ mice tested in both ears, 26% (7/27) had normal hearing in both ears, 33% (9/27) had unilateral hearing impairment (6 in right ear, 3 in left ear), and 41% (11/27) had bilateral hearing impairment (Figure 2D). There were no gender differences in hearing thresholds for either *Df1/+* and WT mice (data not shown; see [20].

We adopted individual ears rather than mice as the unit for analysis after confirming that this approach was reasonable using generalized linear models and randomization tests. Specifically, we used GLMs to investigate how our AEP measures of cortical activity in each mouse, measured over auditory cortex in each hemisphere during monaural stimulation of the contralateral ear, depended on the hearing threshold of the contralateral ear, the hearing threshold of the ipsilateral ear, and the genotype, gender and age of the mouse. Results of the GLM analyses (Table 1) indicated that the only consistently predictive factors were the threshold of the contralateral ear and the genotype of the mouse; the gender of the mouse was also predictive but for only one of the AEP measures analyzed. We then used randomization tests to demonstrate that correlation between paired AEP measures from stimulation of different ears in the same animal was not significantly different from correlation between independent AEP measures from different animals (Supplementary Figure 1). Within-animal correlation fell within the 95% confidence interval for between-animal correlation for all AEP measures and for both P1-N1 and N1-P2 waves. We therefore performed further analyses of our cortical AEP measures grouping AEP recordings by the threshold of the contralateral ear and the genotype of the mouse, with supplemental analysis of gender differences where indicated by the GLM results.

**Table 1.** Generalized linear model (GLM) analyses of factors affecting AEP measures. For each of the AEP measures indicated in rows, a GLM analysis was performed to assess the influence of animal-specific factors. Each mouse contributed two AEP recordings to each GLM analysis, one for each auditory cortical hemisphere and the contralateral, monaurally stimulated ear. Genotype and threshold of the contralateral ear emerged as key factors with significant influence on multiple AEP measures. Gender was a significant factor only for the P1-N1 LDAEP measure. There was no significant influence of the threshold of the ipsilateral ear on any of the AEP measures. Therefore, in further analyses of each AEP measure, we treated AEP recordings corresponding to contralateral ears rather than mice as the unit of analysis.

|  | Genotype | Age | Gender | Threshold of contralateral ear | Threshold of ipsilateral ear |
| --- | --- | --- | --- | --- | --- |
| **P1-N1 amplitude** | <0.001 | 0.698 | 0.632 | 0.015 | 0.151 |
| **N1-P2 amplitude** | 0.005 | 0.726 | 0.894 | 0.182 | 0.107 |
| **P1-N1 central gain** | 0.045 | 0.952 | 0.463 | 0.001 | 0.215 |
| **N1-P2 central gain** | 0.139 | 0.750 | 0.424 | <0.001 | 0.236 |
| **P1-N1 LDAEP** | 0.003 | 0.634 | 0.001 | 0.179 | 0.362 |
| **N1-P2 LDAEP** | 0.115 | 0.429 | 0.059 | 0.209 | 0.229 |
| **P1-N1 TDAEP** | <0.001 | 0.852 | 0.978 | 0.037 | 0.709 |
| **N1-P2 TDAEP** | 0.001 | 0.133 | 0.294 | 0.035 | 0.706 |

### Tone-evoked auditory cortical potentials and brainstem responses confirm increased central auditory gain in *Df1*/+ mice with hearing impairment

Central auditory gain (excitability) can be quantified by comparing the amplitude of central auditory evoked potentials to the amplitude of ABR wave 1, which reflects auditory nerve input to the brain [31]. We recorded tone-evoked cortical AEPs simultaneously with ABRs to test the hypothesis that central auditory gain is elevated in *Df1/+* mice with hearing impairment. A previous study had concluded that *Df1/+* mice with hearing impairment exhibited increased central auditory gain, based on the observation that the ratio of click-evoked AEP to ABR magnitude was abnormally elevated for *Df1/+* ears with hearing impairment [22]. This conclusion is consistent with substantial literature demonstrating that hearing impairment increases excitability of neurons throughout the ascending central auditory system [32]. However, it is possible that alterations in click-evoked AEP/ABR ratio with hearing impairment could arise simply from the different frequency sensitivities of the ABR versus AEP generators (for example, if hearing impairment primarily affected high sound frequencies, which drive the ABR more strongly than the AEP), not from changes in central auditory gain.

To rule out this possible alternative explanation, we investigated the relationship between hearing impairment and the AEP/ABR ratio measure of central auditory gain using 80 dB SPL, 16 kHz tones instead of clicks, and confirmed that elevated central auditory gain is evident in tone-evoked brain activity in *Df1/+* mice. Importantly for this analysis, clicks were used only for categorizing *Df1/+* ears as normal hearing or hearing impaired based on click-evoked ABR threshold, not for analysis of central auditory gain. As expected given the click-evoked ABR thresholds, the amplitude of ABR wave 1 evoked by a 80 dB SPL, 16 kHz tone was much smaller for *Df1/*+ HI ears than for *Df1*/+ NH or WT ears (Figure 3A and C; Kruskal-Wallis test, p < 0.001, post-hoc tests, p*_Df1/_*_+ NH-*Df1/*+ HI_ < 0.001, p*_Df1/_*_+ NH-WT_ = 0.98, p*_Df1/_*_+HI-WT_ < 0.001). However, there was no corresponding drop in the amplitude of cortical AEP waveforms evoked at *Df1/+* HI ears compared to WT ears, and tone stimulation of *Df1/+* NH ears produced AEPs with *larger* P1-N1 amplitude than for WT ears (Figure 3B, D and E; Kruskal-Wallis test, P1-N1, p = 0.0015, post-hoc tests, p*_Df1/_*_+ NH-*Df1/*+ HI_ = 0.017, p*_Df1/_*_+ NH-WT_ = 0.017, p*_Df1/_*_+ HI-WT_ = 0.79; N1-P2, p = 0.034, post-hoc tests, p*_Df1/_*_+ NH-*Df1/*+ HI_ = 0.20, p*_Df1/_*_+ NH-WT_ = 0.20, p*_Df1/_*_+ HI-WT_ = 0.83). The gain of tone-evoked auditory brain activity was elevated in *Df1/+* mice, not only during stimulation of *Df1/+* HI ears but also (for the P1-N1 complex) during stimulation of *Df1/+* NH ears (Figure 3F and G; Kruskal-Wallis tests, P1-N1: p < 0.001, post-hoc tests, p*_Df1/_*_+ NH-*Df1/*+ HI_ = 0.083, p*_Df1/_*_+ NH-WT_ = 0.043, p*_Df1/_*_+ HI-WT_ < 0.001; N1-P2, p < 0.001, post-hoc tests, p*_Df1/_*_+ NH-*Df1/*+ HI_ = 0.037, p*_Df1/_*_+ NH-WT_ = 0.11, p*_Df1/_*_+ HI-WT_ < 0.001).

**Figure 3.**
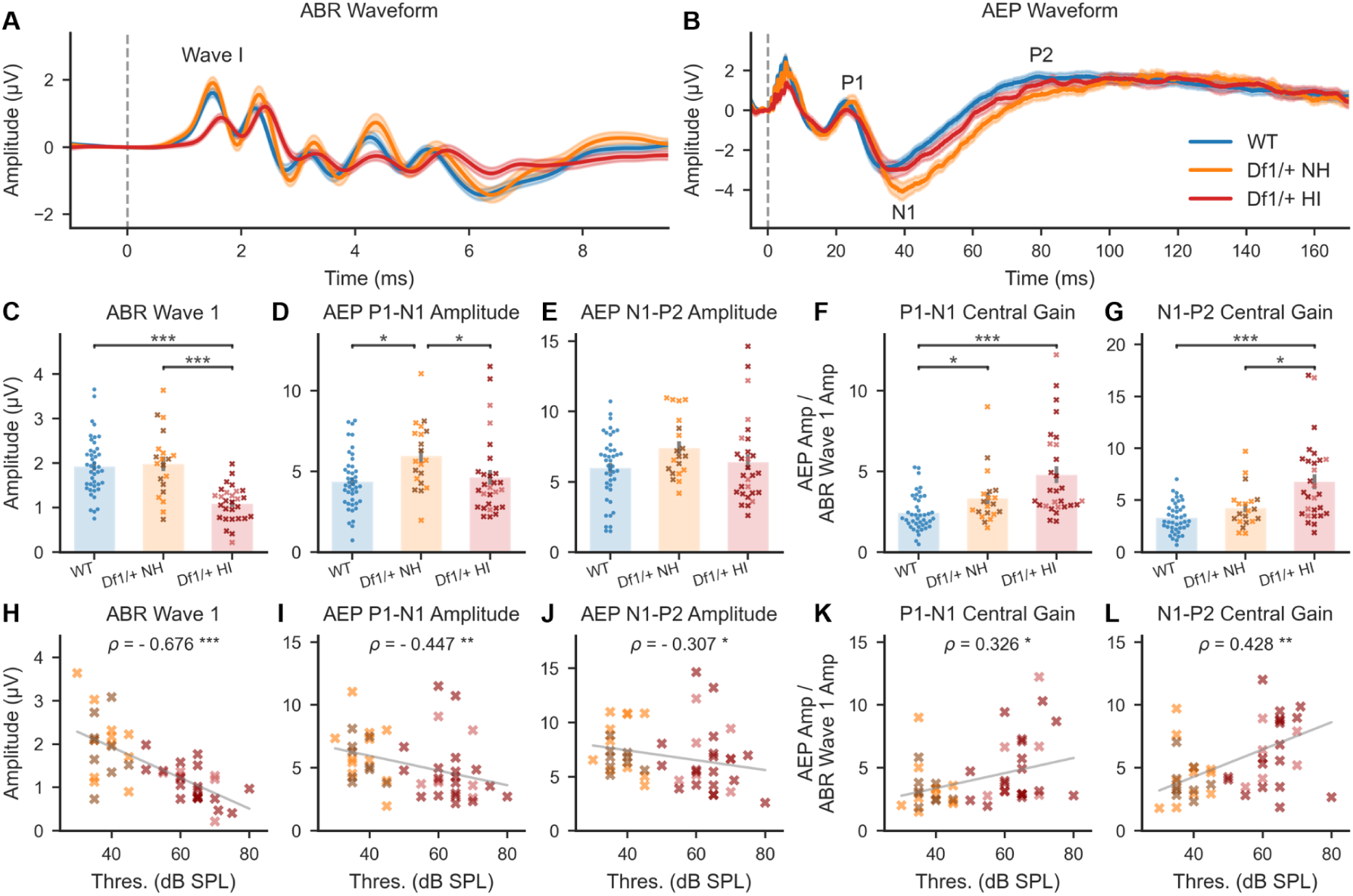
Central auditory gain (AEP/ABR amplitude ratio) was elevated in *Df1/*+ mice for 16 kHz tone stimulus, and positively correlated with click-evoked ABR threshold. (A-B) Averaged tone-evoked ABR waveforms were measured between tested ear bulla and vertex (A) and AEP waveforms were measured from the contralateral hemisphere (B) during monaural stimulation of WT NH, *Df1/*+ NH, and *Df1/*+ HI ears with an 80 dB SPL, 16 kHz tone. Dashed vertical line indicates stimulus onset. (C) Tone-evoked amplitude of ABR wave 1 was significantly reduced for *Df1/+* HI ears versus *Df1/+* NH or WT NH ears. (D-E) However, cortical AEP waves measured contralateral to the stimulated ear showed a different pattern. Amplitude of the AEP P1-N1 complex was largest for stimulation of *Df1/+* NH ears (D); a similar trend was evident for the AEP N1-P2 complex (E). (F-G) Central gain (tone-evoked contralateral AEP amplitude normalized by tone-evoked ABR wave 1 amplitude) was abnormally elevated for the P1-N1 complex during stimulation of *Df1/+* NH ears (F), and for both the P1-N1 and N1-P2 complexes during stimulation of *Df1/+* HI ears (F,G). (H-J) Amplitude of ipsilateral ABR wave 1 (H), contralateral AEP P1-N1 complex (I), or contralateral AEP N1-P2 complex (J) evoked by a 80 dB SPL, 16 kHz tone, versus click-evoked ABR threshold for the stimulated ear. (H) Negative correlation with hearing threshold in the stimulated ear was weaker for tone-evoked AEP than ABR. (K-L) Central gain of tone-evoked responses was positively correlated with click-evoked ABR threshold for both the P1-N1 complex (K) and N1-P2 complex (L). Each animal typically contributed two data points, one for each ear, with NH or HI status determined separately for each ear based on click-evoked ABR thresholds. Hearing sensitivity in the contralateral ear is indicated by the darkness of symbols: darker symbols indicate cases in which the contralateral ear was hearing impaired. Plot conventions: blue circles, WT; yellow cross, *Df1/*+ NH with normal hearing in contralateral ear; brown cross, *Df1/*+ NH with hearing impairment in contralateral ear; red cross, *Df1/*+ HI with normal hearing in contralateral ear; dark red cross, *Df1/*+ HI with hearing impairment in contralateral ear; gray solid line, 2D least-mean-squares best-fit line to the *Df1/*+ data; asterisks, significance threshold for Dunn’s post-hoc tests (C-G) or Spearman’s rank correlation tests (H-L) (*p<0.05, **p<0.01, ***p<0.001).

Further analysis demonstrated that central auditory gain in *Df1/+* mice was positively correlated with the degree of hearing impairment in the stimulated ear. We examined the relationship between click-evoked hearing thresholds and tone-evoked ABR wave 1 amplitude, AEP P1-N1 or N1-P2 amplitude, or AEP/ABR ratio measures of central auditory gain in *Df1/+* mice. Unsurprisingly, the amplitude of tone-evoked ABR wave 1 was negatively correlated with click-evoked hearing threshold in the stimulated ear (Figure 3H; Spearman’s rank correlation test, ρ = -0.68, p < 0.001). Amplitudes of tone-evoked P1-N1 and N1-P2 complexes measured over the contralateral auditory cortex were also negatively correlated with click-evoked hearing thresholds (Figure 3H-I; P1-N1 complex: ρ = -0.45, p = 0.0015; N1-P2 complex: ρ = -0.31, p = 0.034). However, there were robust positive correlations between tone-evoked central auditory gain measures (AEP/ABR ratios) and click-evoked hearing thresholds, for both the P1-N1 and N1-P2 complex (Figure 3K-L; P1-N1 complex AEP/ABR: ρ = 0.33, p = 0.024; N1-P2 complex AEP/ABR: ρ = 0.43, p = 0.0024).

Interestingly, we also found that latencies of late tone-evoked AEP waves were longer in *Df1/+* than WT mice, regardless of hearing impairment (Supplementary Figure 2). Tone stimulation of *Df1/*+ ears with or without hearing impairment evoked N1 and P2 waves with significantly longer latencies than those evoked in WT mice, while P1 wave latencies were not significantly different (Supplementary Figure 2A-C). There was no significant correlation between AEP wave latencies in *Df1/+* mice and the click-evoked hearing threshold in the stimulated ear for any of the three AEP waves (Supplementary Figure 2D-F). Thus, tone-evoked N1 and P2 wave latencies were significantly longer for *Df1/+* than WT mice, even when the stimulated *Df1/+* ears had normal hearing thresholds.

### Higher level-dependent AEP growth in *Df1*/+ mice, with or without hearing impairment

Another measure commonly used to investigate changes in central auditory excitability is growth of cortical AEP amplitude with increasing sound level. A previous study reported steeper level-dependent AEP growth in the *Df(h22q11)/+* mouse model of 22q11.2DS than in WT mice during stimulation with white noise bursts [24]. Here, we used 16 kHz tones to compare level-dependent AEP (LDAEP) growth functions not only between *Df1/+* mice and their WT littermates but also between *Df1/+* mice with or without hearing impairment in the stimulated ear.

We found that tone-evoked cortical AEP wave amplitudes grew more steeply with increasing sound level in *Df1/*+ mice than WT mice, regardless of whether the stimulated *Df1/+* ear was normal hearing or hearing impaired (Figure 4). This effect was clearly evident both in example AEP waveform recordings and in group averages (Figure 4A-B). We calculated the AEP amplitude as a linear function of sound level and used the function slope to represent the level-dependence of the AEP (Figure 4C and E). For both the P1-N1 and N1-P2 complexes, the LDAEP amplitude change was larger in *Df1/+* mice than in their WT littermates (Figure 4D and F; P1-N1: Kruskal-Wallis test, p < 0.001, post-hoc tests, p*_Df1/_*_+NH-*Df1/*+ HI_ = 0.092, p*_Df1/_*_+ NH-WT_ < 0.001, p*_Df1/_*_+ HI-WT_ = 0.040; N1-P2: Kruskal-Wallis test, p = 0.0020, post-hoc tests, p*_Df1/_*_+ NH-*Df1/*+ HI_ = 0.75, p*_Df1/_*_+ NH-WT_ = 0.022, p*_Df1/_*_+ HI-WT_ = 0.0042). P1-N1 LDAEP was larger overall in female than male mice, but the significant effect of genotype on P1-N1 LDAEP was preserved in GLM analysis (Table 1) and also evident as a trend in comparisons between genotypes within gender (Mann-Whitney U rank test, p_male WT-*Df1/+*_=0.034, p_femaleWT-*Df1/+*_=0.061). There were no significant differences in LDAEP amplitude change for data obtained during stimulation of *Df1/+* NH versus *Df1/+* HI ears, suggesting that level-dependent AEP amplitude change could be a biomarker for central auditory abnormalities in *Df1/+* mice that are robust to hearing impairment.

**Figure 4.**
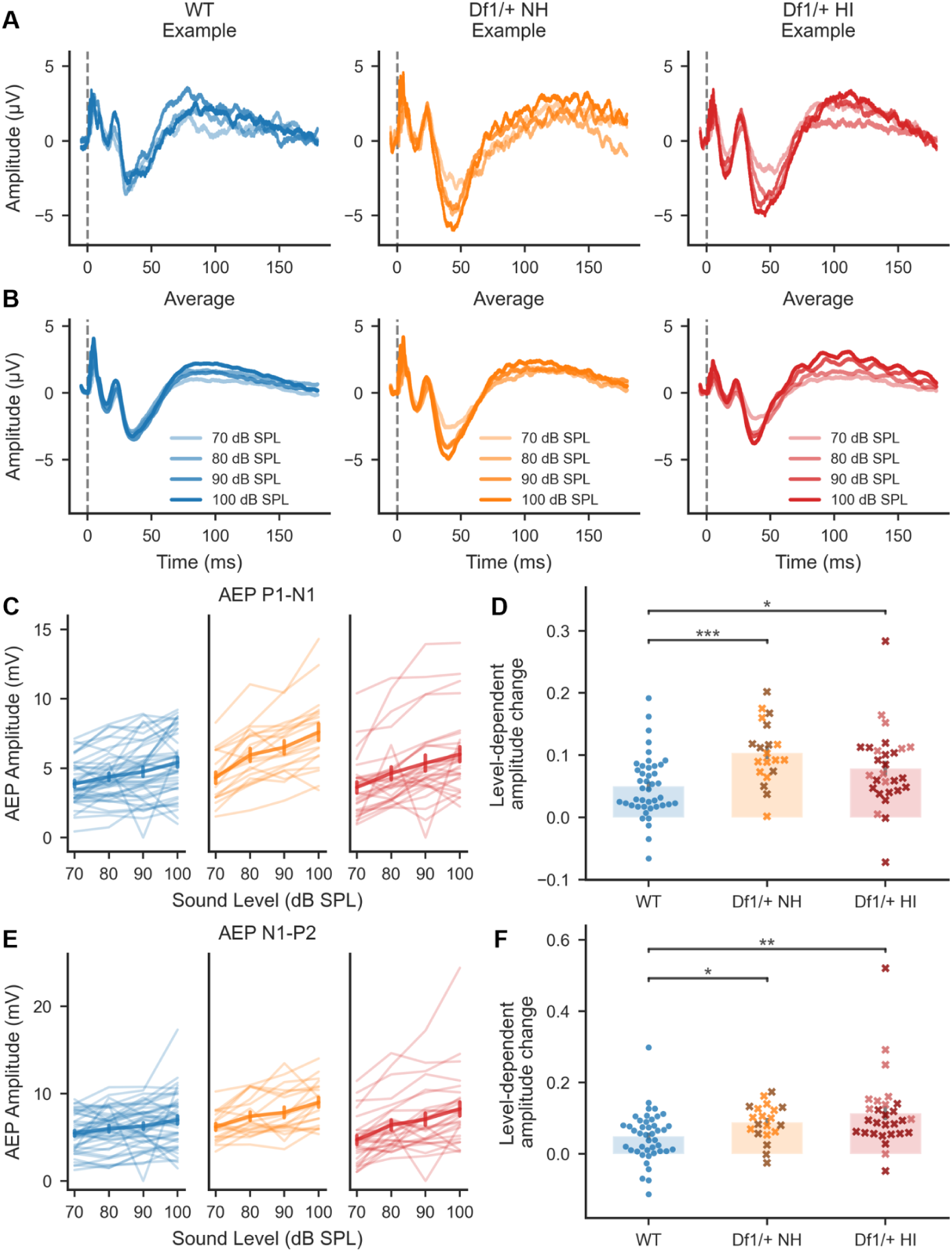
Growth of AEP amplitude with increasing tone intensity was steeper in *Df1/+* mice than WT mice, even for stimulation of *Df1/+* ears with normal hearing. (A-B) Example AEP waveforms (A) and average AEP waveforms (B) evoked by a 16 kHz tone presented at different sound intensity levels. (C,E) AEP amplitudes for the tone-evoked P1-N1 complex (C) and N1-P2 complex (E) grew more steeply with increasing sound level for *Df1/+* mice than WT mice, regardless of the hearing status for the stimulated *Df1/+* ear. Thin lines represent recordings from contralateral hemispheres of individual ears; thick line with error bars shows mean ± SEM. (D,F) For both the P1-N1 complex (D) and the N1-P2 complex (F), slopes of level-dependent AEP growth functions were higher for stimulation of either *Df1/*+ NH or *Df1/*+ HI ears than for WT NH ears. Plot conventions as in Figure 2 H-L.

Level-dependent changes in AEP wave latencies were less robust as a potential biomarker for abnormalities in *Df1/+* mice (Supplementary Figure 3). AEP P1 and N1 latencies decreased more steeply with increasing tone intensity level in *Df1/+* mice compared to their WT littermates, but effects were not consistently significant for the two waves across *Df1/+* NH and *Df1/+* HI ears. For P2, there were no significant differences between the groups in the relationship between wave latency and sound level.

### Higher inter-stimulus-interval time-dependent AEP growth in *Df1/*+ mice with normal hearing

Sound repetition typically causes suppression of auditory evoked activity, and this suppression is stronger at shorter inter-stimulus intervals. The dependence of cortical AEP amplitude on inter-stimulus interval is therefore a measure of repetition suppression; more generally, it is a potential biomarker for abnormalities in central auditory adaptation. A recent study of auditory processing endophenotypes in 22q11.2DS patients showed that tone-evoked AEP amplitudes were more strongly dependent on inter-tone interval (ITI) in patients without psychotic symptoms than in either patients with psychotic symptoms or neurotypical controls [30]. We wondered if similar abnormalities might be evident in *Df1/+* mice, and if so, how these abnormalities might relate to hearing impairment. We measured the sensitivity of the AEP to changes in the temporal structure of the stimulus (time-dependent AEP, TDAEP) using 16kHz, 80 dB SPL tones presented at varying ITIs.

At first we used ITIs varying from 200 ms to 350 ms (Supplementary Figure 4), and observed that AEP amplitude increased with ITI to a plateau in many cases. The turning point for the plateau in WT and *Df1/*+ HI data was between 300 and 350 ms, while the turning point for *Df1/*+ NH data was not clear and could have been at longer ITIs. Therefore, we extended the range of ITIs to 200-450 ms in further experiments.

We found that TDAEP amplitude change was significantly larger for *Df1/+* NH recordings than for either WT or *Df1/+* HI recordings (Figure 5). To quantify TDAEP amplitude change, we calculated the best-fit linear function relating AEP amplitude to the logarithm of ITI duration (Figure 5C and E; note logarithmic scale of x-axis). We used the slope of this function as our measure of TDAEP amplitude change (Figure 5C-F). For both the P1-N1 and N1-P2 complexes, TDAEP amplitude change was abnormally large in *Df1/+* NH recordings, relative to either WT data or *Df1/+* HI data (Figure 5D and F; P1-N1: Kruskal-Wallis test, p < 0.001, post-hoc tests, p*_Df1/_*_+ NH-*Df1/*+ HI_ = 0.060, p*_Df1/_*_+ NH-WT_ < 0.001, p*_Df1/_*_+ HI-WT_ = 0.069; N1-P2: p = 0.0010, post-hoc tests, p*_Df1/_*_+ NH-*Df1/*+ HI_ = 0.10, p*_Df1/_*_+ NH-WT_ < 0.001, p*_Df1/_*_+ HI-WT_ = 0.10). This result indicates that central auditory adaptation to repeated tones differs between *Df1/+* NH and WT mice, and suggests that hearing impairment in *Df1/+* HI mice may either mask or counteract this abnormality.

**Figure 5.**
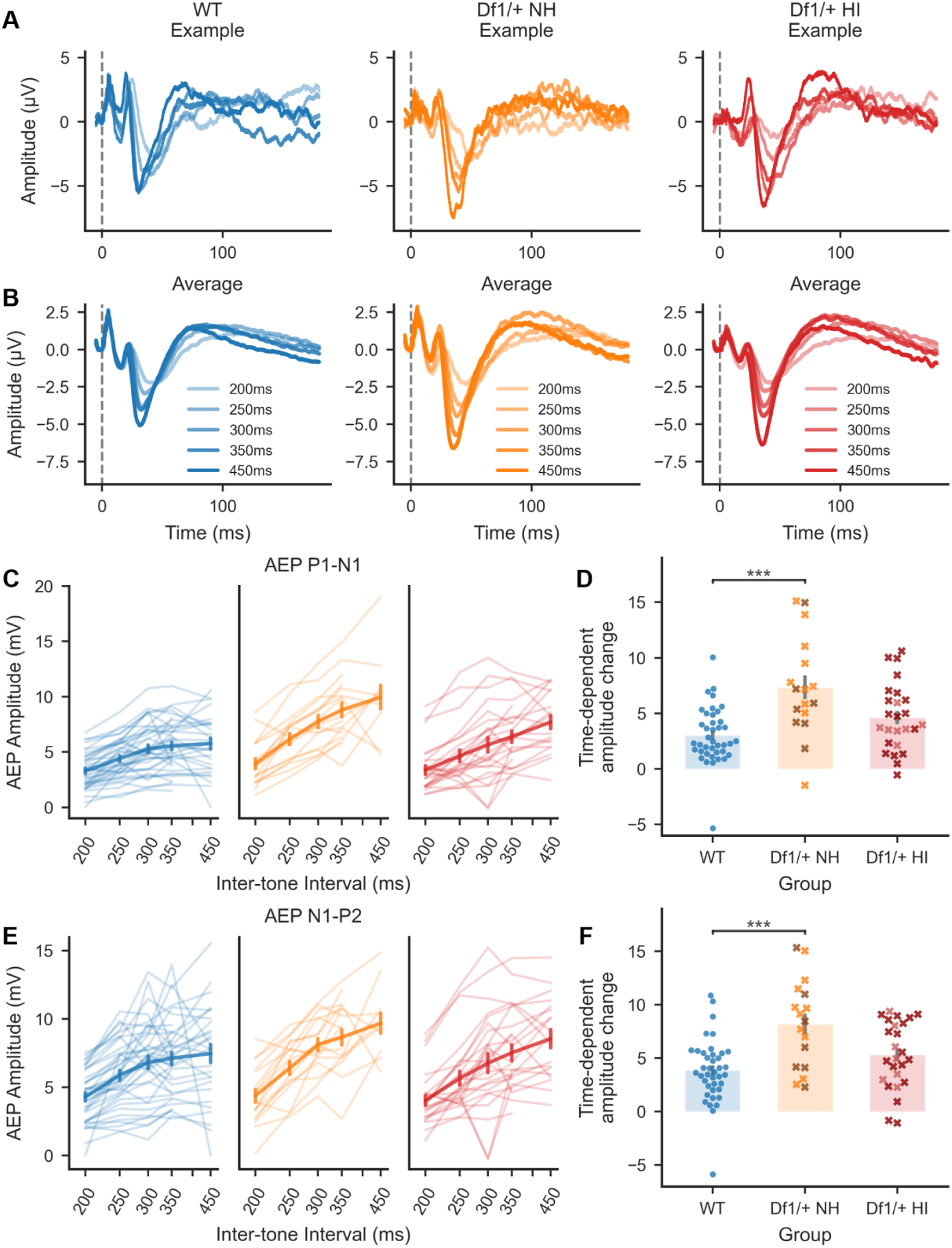
Growth of AEP amplitude with increasing inter-tone interval (ITI) was steeper for stimulation of *Df1/*+ NH ears than for WT ears. (A-B) Example AEP waveforms (A) and average AEP waveforms (B) evoked by 80 dB SPL, 16 kHz tones presented at different ITIs. (C,E) AEP amplitudes for the tone-evoked P1-N1 complex (C) and N1-P2 complex (E) grew more steeply with increasing ITI in *Df1/+* mice than WT mice, particularly at longer ITIs. Note the apparent plateau in AEP amplitude at 300-450 ms ITIs in WT data, and continued growth with ITI in *Df1/+* data, especially for stimulation of *Df1/+* ears with normal hearing. (D,F) For both the P1-N1 complex (D) and the N1-P2 complex (F), slopes of ITI-dependent AEP growth functions were higher for stimulation of *Df1/*+ NH ears than WT ears. A similar trend, but no significant difference, was observed for comparison of ITI-dependent AEP growth-function slopes for stimulation of *Df1/+* HI ears versus WT ears; note however that interpretation of this latter result is complicated by the fact that the fixed-intensity tone stimulus was closer to hearing threshold for *Df1/+* HI ears than WT (NH) ears. Plot conventions as Figure 4.

We also examined the relationship between AEP wave latency and ITI, comparing *Df1/+* NH, *Df1/+* HI, and WT data. For the P1, N1, and P2 waves, latency decreased similarly with increasing ITIs in all groups (Supplementary Figure 5). There were no significant group differences in the relationship between AEP wave latency and ITI for P1 or P2, and for N1, a weak group difference was not robust to post-hoc testing.

## Discussion

Our results reveal multiple abnormalities in cortical auditory evoked potentials (AEPs) in the *Df1/+* mouse model of 22q11.2DS and demonstrate that some of these abnormalities are strongly influenced by mild to moderate hearing impairment, a common co-morbidity of 22q11.2DS. Using one of the best-validated of the surprisingly diverse and unstandardized approaches to mouse AEP measurement in the literature, we measured tone-evoked cortical AEPs along with ABRs in both *Df1/+* mice and their WT littermates. We quantified hearing thresholds ear-by-ear in every mouse using click-evoked ABR thresholds, so that we could exploit the high inter-individual and inter-ear variation in hearing sensitivity typical of *Df1/+* mice (and 22q11.2DS patients) to probe the relationship between AEP abnormalities and hearing impairment.

We found 3 key AEP abnormalities in *Df1/+* mice, which exhibited differing relationships to hearing impairment. Tone-evoked central auditory gain (AEP amplitude normalized by ABR amplitude) was elevated primarily in *Df1/+* mice with hearing impairment, and strongly correlated with the level of hearing impairment in the stimulated ear. Another measure of central auditory excitability --- growth of cortical AEP amplitude with increasing sound level --- was robustly elevated in all *Df1/+* mice, even when hearing was normal in the stimulated ear. Finally, growth of AEP amplitude with increasing inter-tone interval, a measure of central auditory adaptation and recovery from repetition suppression, was abnormal in *Df1/+* mice only when hearing was normal in the tested ear. In summary, these results show that *Df1/+* mice have abnormalities in central auditory gain (AEP/ABR ratio) that are exacerbated by hearing impairment; abnormalities in central auditory excitability (level-dependent AEP growth) that are robust to hearing impairment; and abnormalities in auditory adaptation (inter-stimulus-interval time-dependent AEP growth) that are either counteracted or masked by hearing impairment.

### Implications of results

Two points need to be kept in mind when considering the implications of our results.

First: the unusual nature of hearing impairment in *Df1/+* mice (and 22q11.2DS patients) means that ear-by-ear analysis of hearing sensitivity is essential, and monaural auditory stimulation for AEP measurement is advisable to simplify data analysis and increase statistical power. Like 22q11.2DS patients, *Df1/+* mice are susceptible to chronic middle ear inflammation that can cause mild to moderate hearing impairment in both ears or in only one ear [20,22]. Hearing impairment in our cohort of *Df1/+* mice was frequently unilateral, and there was minimal correlation in hearing thresholds between the two ears. Since auditory cortical AEPs are most strongly driven by contralateral ear stimulation (see for example Postal et al. 2022 [29]), we reasoned that effects of hearing impairment on central auditory function in *Df1/+* mice would best be investigated by recording AEPs over each cortical hemisphere in each animal during monaural stimulation of the contralateral ear. Subsequent GLM analyses showed that our cortical AEP measures depended on the hearing threshold only of the contralateral (stimulated) ear, not the ipsilateral ear. Furthermore, correlation between paired AEP measures from stimulation of different ears in the same animal was not significantly different from correlation between independent AEP measures from different animals. These results simplified further analyses by enabling us to treat AEP recordings corresponding to stimulated ears rather than mice as the unit of analysis. However, it is likely that AEPs measured during binaural stimulation would depend on hearing thresholds in both ears, increasing the variability of experimental results and the complexity of data analysis.

Therefore, one implication of this work in *Df1/+* mice is that AEP abnormalities in 22q11.2DS patients might best be identified using monaural auditory stimulation paradigms, rather than the usual binaural stimulation paradigms (e.g., [13,30,33,34]).

Second: high inter-individual and inter-ear variation in hearing sensitivity among *Df1/+* mice (and 22q11.2DS patients) complicates interpretation of AEP abnormalities. As in most AEP studies of 22q11.2DS patients [13,30,33,34], we presented exactly the same set of auditory stimuli to each subject (and each ear). We did not adjust the sound level of the stimuli to be constant relative to the hearing threshold of the stimulated ear, to avoid the use of extremely loud stimuli for stimulation of ears with moderate hearing impairment (which would have undermined our ability to ensure monaural stimulation using an earplug in the opposite ear). However, the use of fixed-intensity stimuli meant that inevitably, weaker auditory nerve signals were evoked in ears with hearing impairment than in ears with normal hearing. The AEP/ABR ratio measure of central auditory gain attempts to normalize for differences in auditory nerve activity during stimulation of *Df1/+* HI versus *Df1/+* NH or WT ears, making comparisons across all three groups more valid. However, differences in auditory nerve activity are not taken into account by measures of level-dependent AEP growth (LDAEP) and inter-stimulus-interval time-dependent AEP growth (TDAEP), complicating comparisons between hearing-impaired and normal-hearing data groups. Therefore, the most compelling abnormalities in LDAEP and TDAEP are those revealed in *Df1/+* NH versus WT comparisons: higher LDAEP growth, and higher TDAEP growth.

### Biomarkers for central auditory processing abnormalities in 22q11.2DS

Interestingly, similar LDEAP and TDAEP abnormalities have been mentioned previously in studies of 22q11.2DS models or patients. Didriksen et al. (2017) reported higher LDAEP growth in the *Df(h22q11)/+* mouse model of 22q11.2DS than in WT littermates. Notably, hearing thresholds were assessed ear-by-ear using ABR in a separate cohort of these mice and were not significantly different between *Df(h22q11)/+* and WT animals, although unusually high for WT mice [24]. Francisco et al. (2020) [30] found higher TDAEP growth in 22q11.2DS patients without psychotic symptoms compared to neurotypical age-matched controls (although no significant differences were observed between patients with psychotic symptoms and controls). Participants were included in that study only if they passed an audiometric test to ensure that hearing thresholds were no more than 25 dB above normal in both ears. Together with our own data, these two studies suggest that both LDAEP growth and TDAEP growth are abnormally elevated in 22q11.2DS, at least in the absence of hearing impairment. Our work further demonstrates that elevated LDAEP growth may be a particularly useful biomarker for central auditory processing abnormalities in 22q11.2DS, because this AEP abnormality is robust to hearing impairment in the stimulated ear.

### Mechanisms contributing to AEP abnormalities

What mechanisms might underlie the AEP abnormalities we observed in *Df1/+* mice? From a systems-level perspective, increased central auditory gain and LDAEP growth indicate increased excitability in subcortical auditory structures and/or the auditory cortex. The TDAEP results reveal normal adaptation at <250 ms inter-tone intervals in *Df1/+* mice but reduced adaptation (larger AEP) at >300 ms inter-tone intervals, perhaps reflecting normal subcortical adaptation processes but abnormally weak damping of recurrent excitation in higher auditory cortex. Hearing impairment is already known to cause increased excitability and decreased inhibitory synaptic transmission in the auditory cortex [32,35–38]. Moreover, previous work has found that parvalbumin-expressing (PV) inhibitory interneurons, which play a critical role in auditory cortical gain control, adaptation and recurrent excitation [39], are abnormally sparse in the auditory cortex of *Df1/+* mice --- and this PV interneuron deficit correlates with degree of hearing impairment [22]. More generally, deficits in inhibitory signaling have been reported in the prefrontal cortex and hippocampus in other mouse models of 22q11.2DS [40,41], and human 22q11.2DS cerebral cortical organoids have been found to display increased spontaneous firing [42]. It is possible that both the 22q11.2 deletion and hearing impairment alter auditory cortical excitability and adaptation in similar ways, and that auditory brain abnormalities in 22q11.2DS arise from interactions between genetic vulnerability and central consequences of auditory deafferentation.

Previous studies of LDAEP and TDAEP have argued that abnormalities in these measures are linked to alterations in particular neuromodulator or neurotransmitter systems. LDAEP in particular is often described as a measure of cortical serotonergic activity, based on results from pharmacological studies linking increased level dependence of the N1/P2 waves with decreased serotonergic neurotransmission [43,44]. However, both LDAEP and TDAEP are blunted by administration of N-methyl-D-aspartate (NMDA) receptor blockers [45], indicating that the neuropharmacology of these AEP measures may be complex. Moreover, cortical AEPs are shaped by integration, adaptation, and synchrony of neural population activity in multiple brain areas along the ascending auditory pathway, so it is not necessarily reasonable to expect a simple one-to-one relationship between specific AEP abnormalities and particular neuromodulator or neurotransmitter systems. We suggest that systems-level mechanistic explanations for AEP abnormalities, such as increased central auditory excitability and reduced adaptation at longer time scales, are more likely to be translationally useful.

### Conclusions and future directions

We found multiple abnormalities in cortical auditory evoked potentials (AEPs) in the *Df1/+* mouse model of 22q11.2DS, some of which were strongly influenced by hearing impairment, a common co-morbidity of 22q11.2DS. *Df1/+* mice exhibited 3 key AEP abnormalities: (1) increased central auditory gain, which was especially pronounced for stimulation of ears with hearing impairment; (2) increased growth of AEP amplitude with sound level, regardless of the presence or absence of hearing impairment in the stimulated ear; and (3) increased growth of AEP amplitude with inter-stimulus-interval time duration when hearing was normal in the stimulated ear. Based on these results, we suggest that level-dependent AEP growth may be especially useful as a biomarker for auditory brain abnormalities in 22q11.2DS, because this measure was robust to hearing impairment in the stimulated ear.

More generally, we conclude that auditory deafferentation and genetic risk for 22q11.2DS can have both independent and interactive effects on evoked-potential measures of auditory brain function. Further experiments are needed to address three limitations of the present work. First, a more complete dissociation of effects of genetic risk and hearing sensitivity could be achieved if the group comparisons included WT mice with induced hearing impairment similar in severity and developmental timecourse to that caused by otitis media in *Df1/+* mice. Second, the use of ketamine as the anesthetic could have affected the AEP results by blocking NMDA receptors (Schwender et al. 1993; Teichert et al. 2017); this potential confound might be avoided in future by recording in awake mice. Third, and most importantly, our results were obtained in a mouse model of 22q11.2DS. Future studies in humans will be required to test the hypothesis that co-morbid hearing impairment could be a “second-hit” risk factor for brain abnormalities and psychiatric disease in 22q11.2DS patients.

## Acknowledgements

We thank Dr. Alain de Cheveigne for advising on data analysis and Ashviniy Thamilmaran for assistance with literature data collection. We also thank Dr. Pedro J. Gonçalves for discussions of the comparison between experimental data and predictions from computational models. This research was funded by the UCL Institute for Mental Health (Small Grant 2022 to JFL) and the UK Medical Research Council (MR/P006221/1 to JFL) and also supported by the NIHR UCLH BRC Hearing Health Theme.

## Other information

Confirmation of references for Figure 1, for article proof:

#01 [46]; #02 [47]; #03 [48]; #04 [49]; #05 [50]; #06 [51]; #07 [52]; #08 [53]; #09 [54];

#10 [55]; #11 [56]; #12 [57]; #13 [58]; #14 [59]; #15 [26]; #16 [25]; #17 [24]; #18 [60];

#19 [61]; #20 [27]; #21 [62]; #22 [29]; #23 [63]; #24 [64]; #25 [65]; #26 [66]; #27 [67];

#28 [68]; #29 [69]; #30 [70]; #31 [71]; #32 [72]; #33 [73]; #34 [74]; #35 [75]; #36 [76];

#37 [77]; #38 [78]; #39 [79]; #40 [80]; #41 [81]; #42 [82]; #43 [83];

#44 [84]; #45 [85]; #46 [86]; #47 [87]; #48 [88]; #49 [89]; #50 [90]

## Supplementary Information

### Methods and Materials

#### Acoustic stimulation

Auditory stimuli were generated at a sample rate of 195,312.5 Hz using a digital signal processor (Tucker Davis Technologies, TDT RX6), attenuated as needed (TDT PA5), amplified (TDT SA1), and presented using a free-field speaker (TDT FF1) positioned 10 cm from the ear directed toward the speaker. Speaker output was calibrated to within 1.5 dB of target values before each set of experiments using a G.R.A.S. free-field 1/4" microphone, placed at the location of the ear to be tested.

#### Experimental procedures

Animals were anesthetized with intraperitoneally injected ketamine (70-100 mg/kg) and medetomidine (0.24-0.50 mg/kg) for recording, and were also given subcutaneous doses of carprofen (4.8-8.3 mg/kg) for analgesia and atropine (0.10-0.16 mg/kg) to minimize respiratory secretions. Supplementary doses of ketamine or ketamine/medetomidine mixture were administered if a toe-pinch response or whisker twitching was observed. Body temperature was maintained at 37-38°C using a homeothermic blanket (Harvard Apparatus). Each mouse was placed on an elevated platform and oriented with the tested ear directed toward the speaker. The opposite ear was blocked with an earplug during the recording to ensure monaural stimulation.

Subdermal electrodes (Rochester Medical) were inserted at the vertex; the bulla of the tested ear; the ipsilateral and contralateral auditory cortex relative to the tested ear; and the olfactory bulb (ground electrode). Data was acquired at a 24,414 Hz sample rate (TDT RX5) using a low-impedance headstage and signal amplifier (TDT RA4LI and RA16SD, 20x gain overall, 2.2 Hz - 7.5 kHz filtering) along with a custom low-pass filter designed to remove attenuation switching transients (100 kHz cutoff). Stimulus presentation and data acquisition was controlled using software from TDT (Brainware) and custom software written in MATLAB.

#### Data pre-processing

To check the quality of AEP recordings, distributions of root mean square (RMS) values of the 1000 trials in every AEP recording session were checked. No clear outlier trials that were discrete from the rest of RMS distribution were identified for any of the recordings; therefore, we didn’t exclude any trials from the analysis. Three key deflections of the AEP waveform (P1, N1, and P2) were manually selected from the averaged waveform of all 1000 trials after removing heartbeat noise [1]. Heartbeats were identified as large differences between the detrended waveform and a detrended, smoothed waveform, and eliminated by subtracting these differences from the detrended waveform.

Across all recordings from a stimulated ear and the contralateral auditory cortex, 4 recordings were excluded because no clear P1, N1 and P2 could be identified, and 2 were excluded because the mice died from possible anaesthetic overdose during recording.

## Supplementary Figures

**Supplementary Figure 1.**
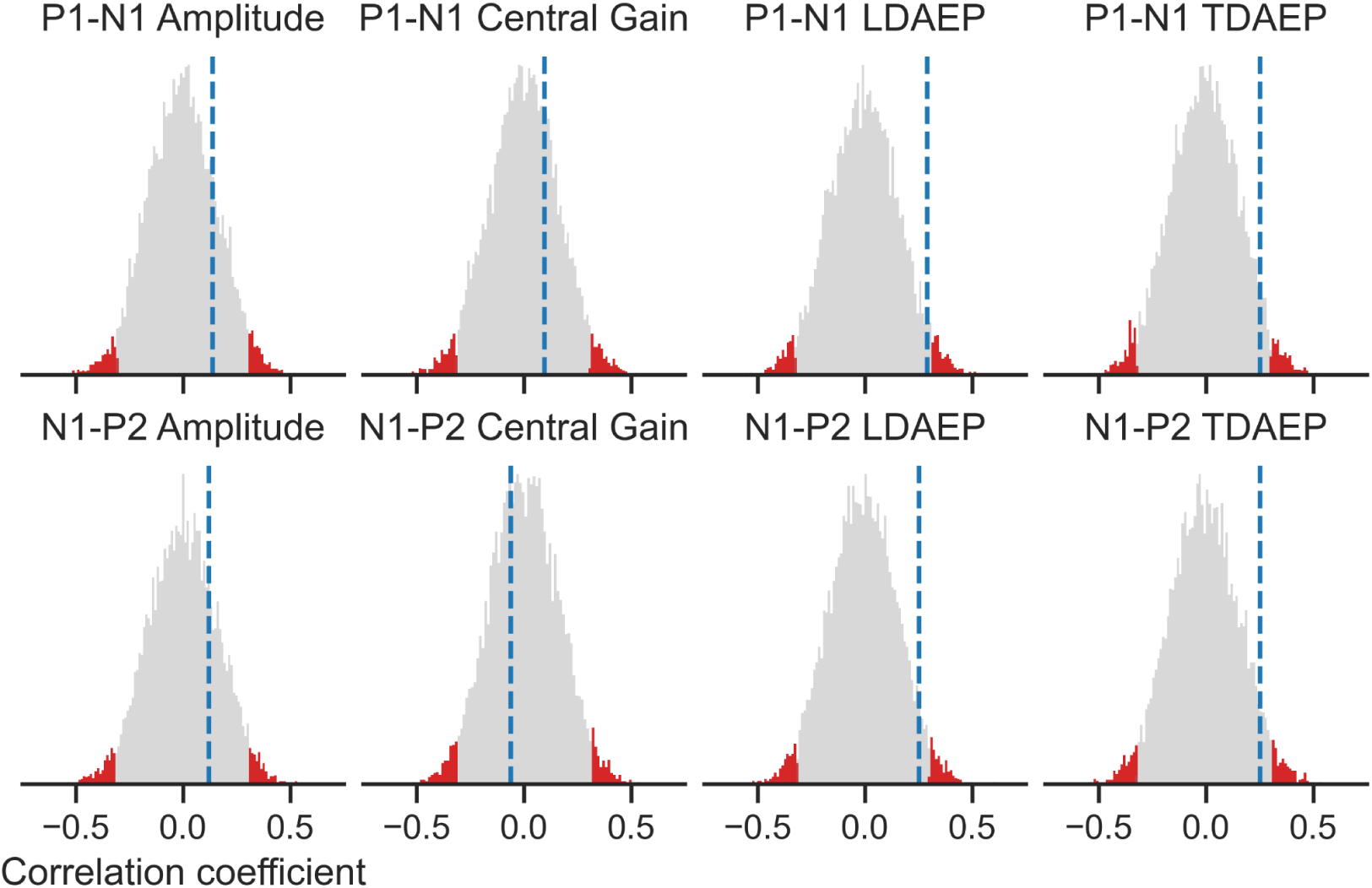
Correlation between paired AEP measures from stimulation of different ears in the same animal was not significantly different from correlation between independent AEP measures from different animals. For most animals, we had two sets of AEP measures, obtained from recordings over the contralateral auditory cortex during monaural stimulation of each ear. To determine whether correlation between paired AEP measures was significantly different from that expected for independent AEP measures, we calculated Spearman’s rank correlation coefficient for the paired data (blue dashed vertical line in each plot) and compared it to the distribution of correlation coefficients obtained in 10,000 randomizations of the pairing to different animals (gray histograms; lowest 2.5% and highest 2.5% of correlation coefficients highlighted in red). Within-animal correlation in AEP measures fell within the 95% confidence interval for between-animal correlation for both P1-N1 and N1-P2 waves (rows) and for all AEP measures (columns).

**Supplementary Figure 2.**
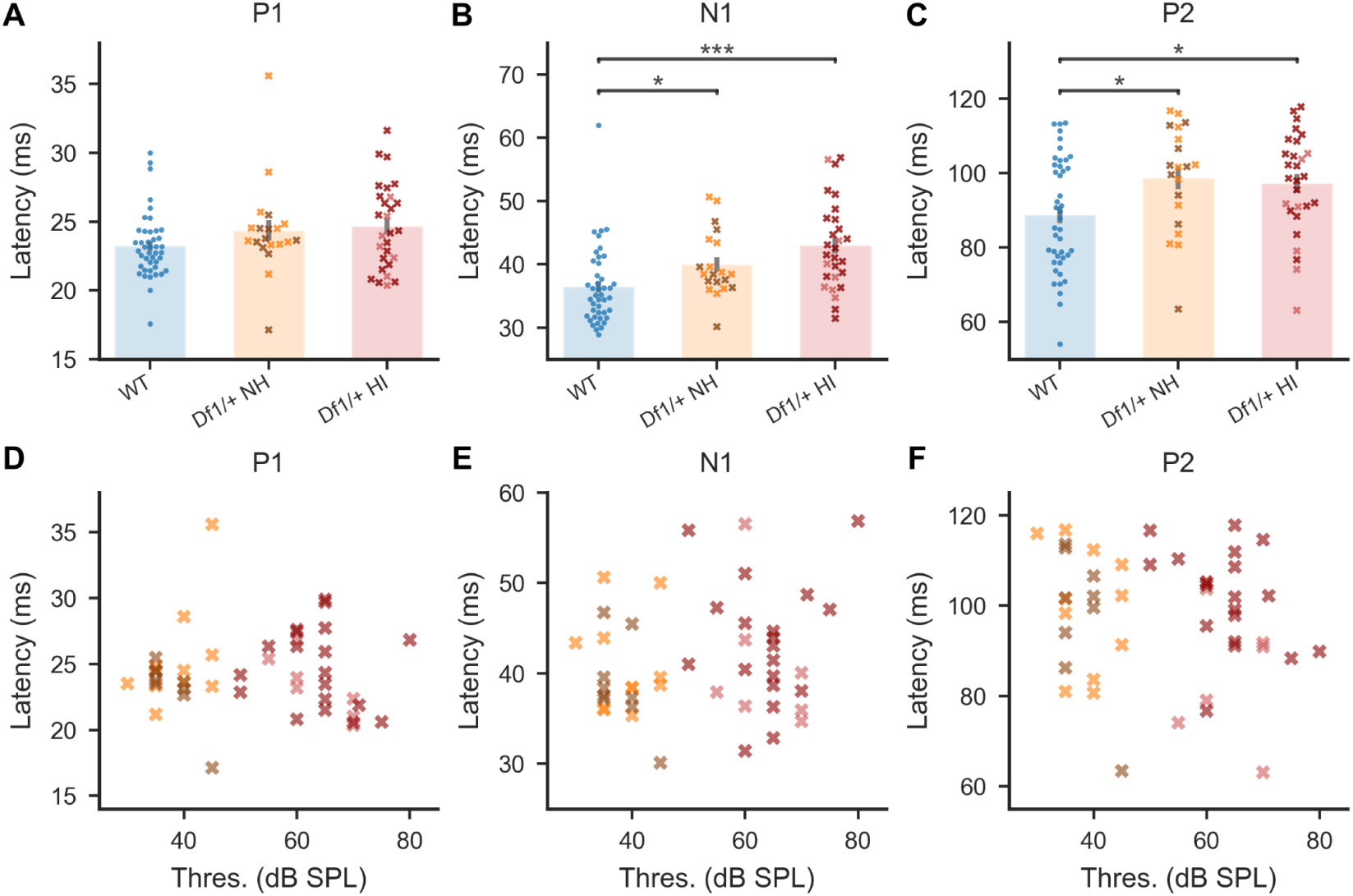
Latencies of AEP waves N1 and P2 evoked by a fixed-intensity tone were abnormally elevated in *Df1*/+ mice and uncorrelated with click-evoked ABR threshold in the stimulated ear. (A-C) A fixed-intensity tone (80 dB SPL, 16 kHz) evoked longer latency N1 and P2 waves in *Df1/+* mice than WT mice, regardless of whether the stimulated *Df1/+* ear had a normal or impaired hearing threshold (B,C; N1: *Df1/*+ NH 39.96 ± 5.05 ms, *Df1/*+ HI 42.98 ± 6.82 ms, WT 36.39 ± 6.04 ms, p < 0.001, post-hoc tests, p*_Df1/_*_+ NH-*Df1/*+ HI_ = 0.23, p*_Df1/_*_+ NH-WT_ = 0.020, p*_Df1/_*_+ HI-WT_ < 0.001; P2:, *Df1/*+ NH 98.64 ± 13.75 ms, *Df1/*+ HI 97.25 ± 13.28 ms, WT 88.74 ± 15.09 ms, p = 0.015, post-hoc tests, p*_Df1/_*_+ NH-*Df1/*+ HI_ = 0.70, p*_Df1/_*_+ NH-WT_ = 0.042, p*_Df1/_*_+ HI-WT_ = 0.046). There were no significant differences between WT and *Df1/+* data for latency of the P1 wave (A; P1: *Df1/*+ NH 24.35 ± 3.31 ms, *Df1/*+ HI 24.66 ± 3.05 ms, WT 23.24 ± 2.33 ms, Kruskal-Wallis test, p = 0.057). (D-F) No significant correlations between the latency of AEP waves evoked by an 80 dB SPL, 16 kHz tone and the click-evoked ABR threshold of the stimulated *Df1*/+ ear (Spearman’s rank correlation tests, P1, N1 and P2 all p > 0.05). Plot conventions as in Figure 3.

**Supplementary Figure 3.**
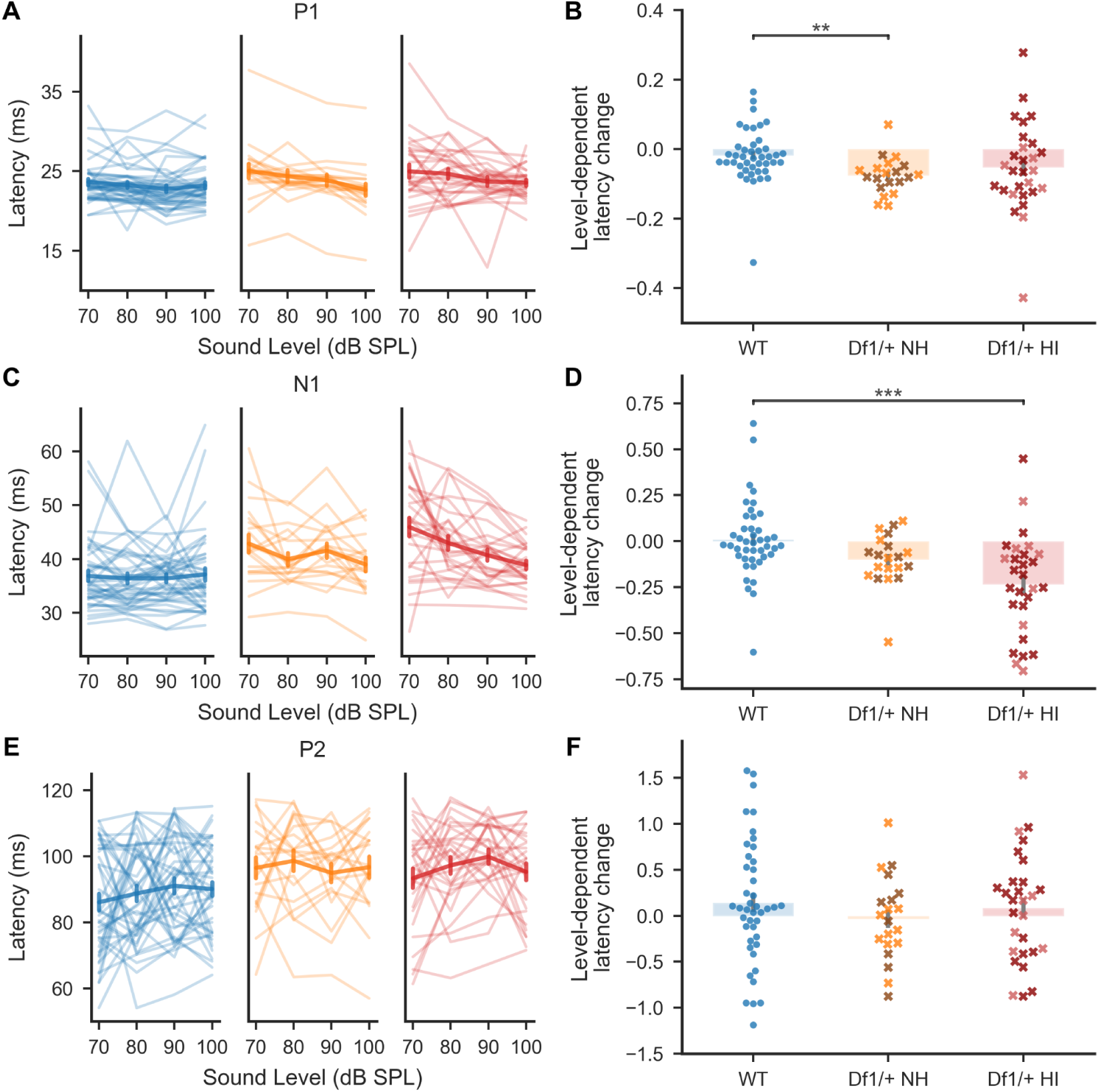
Decrease of AEP wave latency with increasing tone intensity was slightly more pronounced in *Df1/+* mice than WT mice. (A,C,E) Latency for the P1 (A) and N1 (C) AEP waves evoked by a 16 kHz tone dropped slightly with increasing sound level, while latency for the P2 wave (E) was stable across sound levels in all groups (P1: Kruskal-Wallis test, p = 0.0017, post-hoc tests, p*_Df1/_*_+ NH-*Df1/*+ HI_ = 0.18, p*_Df1/_*_+ NH-WT_ = 0.0019, p*_Df1/_*_+ HI-WT_ = 0.059; N1: Kruskal-Wallis test, p < 0.001, post-hoc tests, p*_Df1/_*_+ NH-*Df1/*+ HI_ = 0.090, p*_Df1/_*_+ NH-WT_ = 0.072, p*_Df1/_*_+ HI-WT_ < 0.001; P2: Kruskal-Wallis test, p = 0.61). (B,D,F) Slopes of level-dependent latency growth functions for P1 (B) and N1 (D) were slightly steeper in Df1/+ ears, but the growth function for P2 didn’t show significant between-group differences (F). Plot conventions as in Figure 4.

**Supplementary Figure 4.**
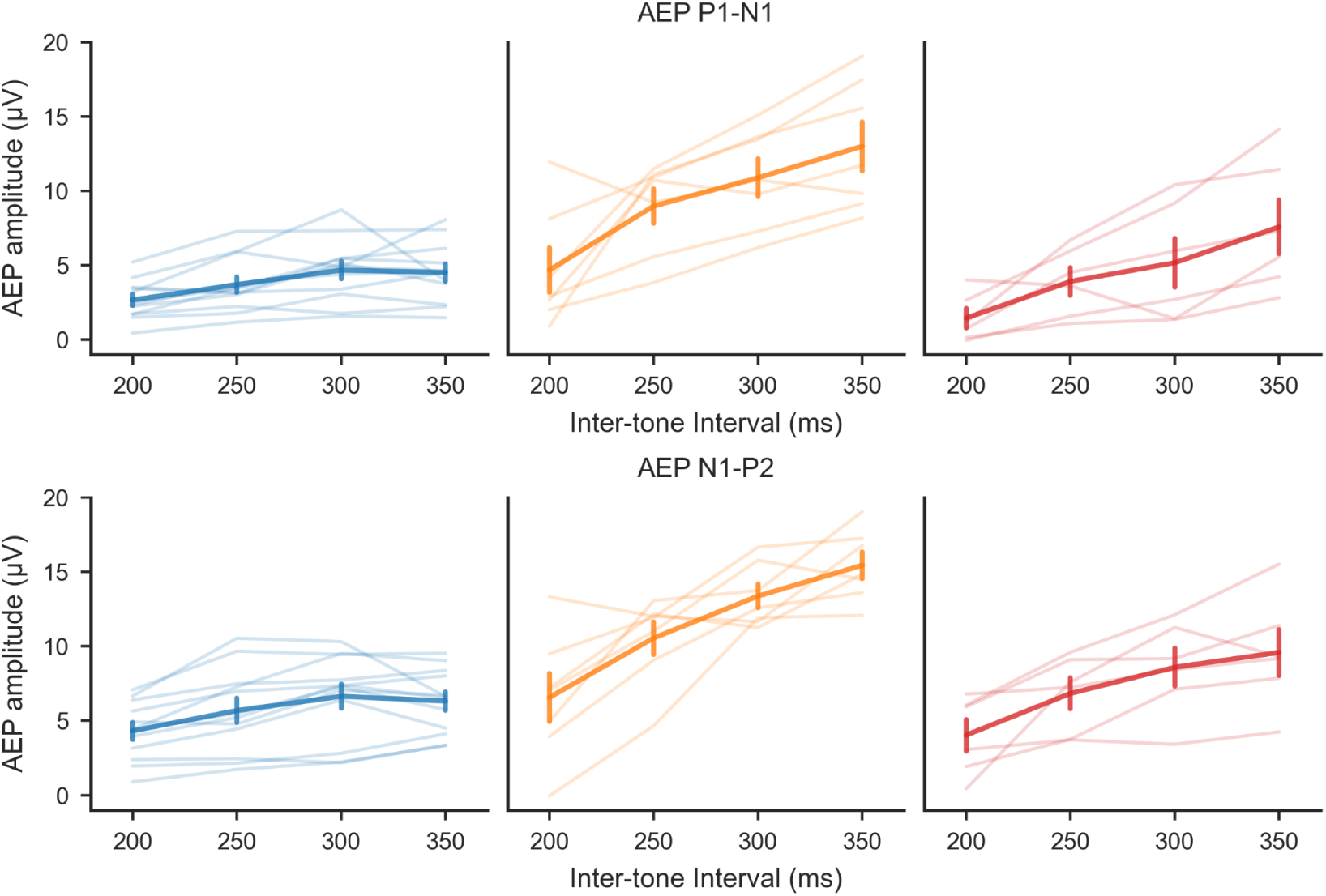
Plateau in growth of AEP wave amplitude at 300-350 ms inter-tone intervals (ITIs) for WT but not *Df1/+* mice. For both P1-N1 and N1-P2 complexes, AEP amplitude showed a plateau at longer ITIs (300-350 ms) in WT mice. In contrast, AEP wave amplitude kept increasing over this ITI interval in *Df1*/+ mice, for stimulation of both *Df1/+* NH and *Df1*/+ HI ears. Solid lines represent AEP recordings contralateral to the stimulated ears; thick line with error bars shows mean ± SEM.

**Supplementary Figure 5.**
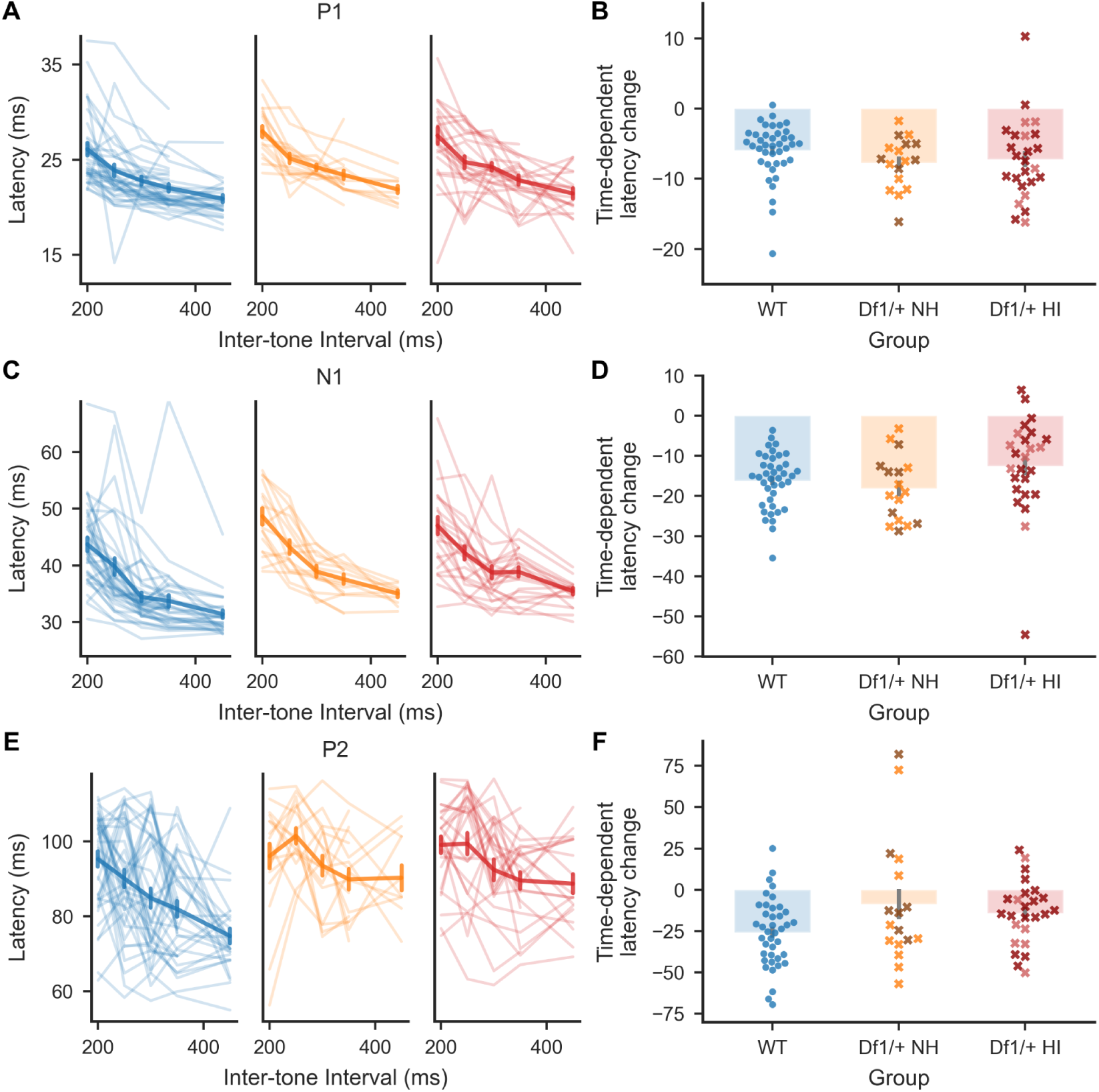
No differences between *Df1/+* and WT mice in the relationship between AEP wave latency and ITI for a fixed-intensity tone. (A,C,E) Latencies of the P1 (A), N1 (C), and P2 (E) waves evoked by an 80 dB SPL, 16 kHz tone dropped with increasing ITI. Note that although AEP wave latencies were slightly elevated overall for both *Df1/+* NH and *Df1/+* HI data compared to WT data (see also Supplementary Figure 1), the relationship between wave latency and ITI was generally consistent across the three groups. Slopes of the ITI-dependent latency growth functions were not significantly different between groups for P1 or P2, and were different for N1 only overall, not in post-hoc tests (Kruskal-Wallis test, P1: p = 0.11; N1: p = 0.040, post-hoc tests, p*_Df1/_*_+ NH-*Df1/*+ HI_ = 0.051, p*_Df1/_*_+ NH-WT_ = 0.39, p*_Df1/_*_+ HI-WT_ = 0.10; P2: p = 0.086). Plot conventions as in Figure 4.

